# A novel approach for combining task-dependent gamma with alpha and beta power modulation for enhanced identification of eloquent cortical areas using ECoG in patients with medical-refractory epilepsy

**DOI:** 10.1101/677195

**Authors:** M.E. Archila-Meléndez, G. Valente, E. Gommer, R.P.W. Rouhl, O.E.M.G. Schijns, J.T.A. Dings, D.M.W. Hilkman, B.M. Jansma, V.H.J.M. van Kranen-Mastenbroek, M.J. Roberts

## Abstract

Electrical stimulation mapping (ESM) is the gold standard for identification of “eloquent” areas prior to resection of epileptogenic tissue, however, it is time consuming and may cause side effects, especially stimulation-induced seizures and after-discharges. Broadband gamma activity (55 – 200 Hz) recorded with subdural electrocorticography (ECoG) during cognitive tasks has been proposed as an attractive tool for mapping cortical areas with specific function but until now has not proven definitive clinical value. Fewer studies have addressed whether the alpha (8 – 12 Hz) and beta (15 – 25 Hz) band activity could also be used to improve eloquent cortex identification. We compared alpha, beta and broadband gamma activity, and their combination for the identification of eloquent cortical areas defined by ESM. Ten patients participated in a delayed-match-to-sample task, where syllable sounds were matched to visually presented letters and responses given by keyboard. We used a generalized linear model (GLM) approach to find the optimal weighting of low frequency bands and broadband gamma power to predict the ESM categories. Broadband gamma activity increased more in eloquent areas than in non-eloquent areas and this difference had a diagnostic ability (area under (AU) the receiving operating characteristic curve - AUROC) of ∼70%. Both alpha and beta power decreased more in eloquent areas. Alpha power had lower AUROC than broadband gamma while beta had similar AUROC. AUROC was enhanced by the combination of alpha and broadband gamma (3% improvement) and by the combination of beta and broadband gamma (7% improvement) over the use of broadband gamma alone. Further analysis showed that the relative performance of broadband gamma and low frequency bands depended on multiple factors including the time period of the cognitive task, the location of the electrodes and the patient’s attention to the stimulus. However, the combination of beta band and broadband gamma always gave the best performance. We show how ECoG power modulation from cognitive testing periods can be used to map the probability of eloquence by ESM and how this probability can be used as an aid for optimal ESM planning. We conclude that low frequency power during cognitive testing can contribute to the identification of eloquent areas in patients with focal refractory epilepsy improving its precision but does not replace the need of ESM.

**Highlights:**

- Gamma, alpha and beta band activity has significant diagnostic ability to identify ESM defined eloquent cortical areas.
- We present a novel method to combine gamma and low frequency activity for enhanced identification.
- We quantify how identification is dependent on analysis time window, cortical function, and patient’s attentional engagement.
- With further development, this approach may offer an alternative to ESM mapping with reduced burden for patients.

## 1. Introduction

Invasive cortical mapping for the precise characterization of ‘eloquent’ cortical areas is necessary to avoid or decrease neurological or cognitive complications following resection of pathologic brain tissue. In the current gold standard, electrical cortical stimulation mapping (ESM), electrical stimulation of subdural electrode-pairs, disrupts function or produces neurological symptoms (Hamberger, 2007; Penfield and Boldrey, 1937). If the stimulation impairs a specific cognitive function (e.g., speech production) or produces neurological symptoms (e.g., paraesthesia), the underlying cortex is labelled as ‘eloquent’ and is preserved during resection. Despite its usefulness (Ojemann et al., 1989), ESM can elicit after-discharges or seizures (Lee et al., 2010; Lesser et al., 1984), or induce pain (Lesser et al., 1985). In addition, it requires the patient’s continuous compliance, rendering it challenging to use especially in paediatric populations (Arya et al., 2015). The procedure is also time consuming, requiring the individual testing of each implanted electrode, restricting the maximum number of electrodes that can be tested and precluding the use of high density arrays (Bouchard et al., 2013; Mesgarani et al., 2014; Muller et al., 2016) so limiting the spatial resolution of the technique (Hermiz et al., 2018). These and other factors motivate the search for alternative ways to identify eloquent cortex (Brunner et al., 2009; Crone et al., 2006, 1998b, 1998a; Lachaux et al., 2007; Vansteensel et al., 2010).

Activation of neuronal networks leads to a change in the spectral power of electrical field potentials of local neuronal populations (Buzsaki, 2004; Buzsáki et al., 2012; Fries et al., 2007). For example, different studies have reported enhancement of gamma band power (>30Hz) during processing of visual (Gray et al., 1989) and auditory (Brosch et al., 2002) stimuli, motor action preparation (Pfurtscheller et al., 1993; Vansteensel et al., 2013), and sensorimotor integration (Murthy and Fetz, 1992). Gamma power varies with high temporal and spatial resolution, such that increasing gamma power is specific to active neuronal populations (Aoki et al., 1999, p. 199; Arya et al., 2018; Buzsáki et al., 2012; Crone et al., 2006, 1998a; Hamilton et al., 2018; Leuthardt et al., 2007; Miller et al., 2007; Nagasawa et al., 2010; Sinai et al., 2005; Wu et al., 2010). Consequently, gamma modulation has been proposed as an alternative for ESM (Aoki et al., 1999; Arya et al., 2017; Brunner et al., 2009; Crone et al., 2006, 1998a; Lachaux et al., 2003; Leuthardt et al., 2007; Miller et al., 2007; Sinai et al., 2005; Vansteensel et al., 2013; Wang et al., 2016; Wu et al., 2010) whereby a task-dependent increase in gamma power indicates eloquent cortex. However, results are mixed and gamma band-based mapping typically has insufficient accuracy in the identification of ESM results, with some studies (Wu et al., 2010) reporting high sensitivity but low specificity, while others report the opposite (Bauer et al., 2013). In a recent review Arya et al. highlighted the heterogeneity in the diagnostic threshold and the cognitive task employed (Arya et al. 2018). In addition there is heterogeneity in the criteria used during ESM (Mooij et al., 2018). Moreover, it has recently been suggested that broadband gamma activity composing cortical field potentials, registered by electrocorticography (ECoG) electrodes, may over-represent dendritic processes more than neuronal action potentials (Leszczynski et al., 2019).

Activation of neuronal networks is typically accompanied by a reduction in power in the alpha (8 – 12 Hz) and beta (15 – 25 Hz) frequency bands. Recent empirical evidence demonstrates that alpha and beta power is related to active inhibition processes (Jensen and Mazaheri, 2010) and can be highly spatially specific to activated populations (de Pesters et al., 2016; Muller et al., 2016). Thus, activity in lower frequency bands represents a plausible source of additional information for cortical mapping. Moreover, given that different frequency bands have different functions (Scheeringa and Fries, 2017) and are not directly correlated to each other (Bonaiuto et al., 2018), additional information may come from the combination of frequency bands. A limited number of studies have investigated the use of lower frequency bands for cortical mapping (Bauer et al., 2013; Crone et al., 1998b; Hermes et al., 2012; Leuthardt et al., 2007; Sinai et al., 2005; Vansteensel et al., 2013; Wu et al., 2010). The results from these studies have been have been variable in terms of mapping performance and the combination of frequency bands has not yet been fully investigated (Leuthardt et al., 2007; Wu et al., 2010). In addition, while there is considerable heterogeneity between studies in the task employed, few studies have directly compared different tasks in the same group of patients. Thus, little is known about how the cognitive engagement of the patient, i.e., the task performed during cortical mapping, impacts the quality of ECoG based mapping.

We hypothesized that alpha and beta band power modulation, either on their own or in combination with the broadband gamma power modulation, could enhance the accuracy of the identification of eloquent cortex compared with the use of broadband gamma alone.

To investigate this, we registered ECoG signals from subdural electrodes in 10 patients with drug resistant focal epilepsy who underwent ESM for the identification of eloquent cortex. Patients performed a delayed match-to-sample (DMTS) task or listened to the same stimuli without an active task. We show that beta band power generally had equal diagnostic ability to gamma, while alpha power was less effective. Withdrawing attention from the stimuli reduced the diagnostic ability. The combination of gamma and beta frequency bands, using a Generalized Linear Model (GLM) was consistently better than either band individually.

## 2. Materials and Methods

### 2.1. Participants

We included 10 patients (mean age: 40.1 years; 4 females; 3 left handed) with drug-resistant focal epilepsy, who underwent continuous ECoG with subdural electrodes as part of pre-surgical evaluation. The remaining patients of our full data set were excluded from this study as ESM was not performed in those patients. Patients volunteered as participants for the study and performed the DMTS task. The study complies with the Declaration of Helsinki for research studies in humans. Written informed consent was obtained from all patients before admittance to the ward. All experimental procedures were approved by the Medical Ethical Committee of the Maastricht University Medical Center, Maastricht, Netherlands.

### 2.2. Electrode implantation

The electrode implantation scheme was strictly chosen according to the individual patient protocol depending on the hypothesis of the probable location of the epileptogenic zone, as established by the multidisciplinary epilepsy surgery-working group of the Academic Center for Epileptology. If clinically indicated, language and verbal memory lateralization was established preoperatively using fMRI or a Wada test (Wada and Rasmussen, 2007). The number of implanted electrodes per patient varied, with electrodes’ contact points ranging between 24 and 88 per patient. Henceforth we will use the word *electrode* to refer to the electrode’s contact point. The day after implantation, a computer tomography and a 3T magnetic resonance imaging scan of the brain was acquired to verify the location of implanted electrodes and to exclude post-surgical complications.

### 2.3. Electrical-cortical stimulation mapping (ESM)

Electrical-cortical stimulation mapping (ESM) was performed using bipolar stimulation between electrode pairs sequentially, selecting neighbouring electrode pairs of the subdural grid and/or strips. The procedure for ESM mapping differs between centres, with consequences for the reproducibility of comparisons between ESM and ECoG response based mapping approaches (Mooij et al., 2018). In our study ESM mapping proceeded as follows: The stimulation of an electrode-grid was planned by selecting non-continuous electrode pairs (Figure 1A). Bipolar electrical stimulation was applied between a pair of electrodes (e.g. electrode pair 1-2). If no neurological symptoms or function disturbances were observed, electrical stimulation was applied to the next planned pair (pair 3-4), which included neither of the electrodes included in the first pair. If instead, symptoms were generated by the stimulation of pair 1-2, then electrical stimulation was applied to a neighbouring pair of electrodes that included one of the previously stimulated electrodes and (if possible) one functionally silent electrode e.g. pair 1-5. If this pair resulted in symptoms (Figure 1B) electrode 1 would be labelled as eloquent. If a pair that includes the second electrode of the first pair (e.g. pair 2-3) resulted in no symptoms, that electrode would be labelled as non-eloquent. In a different scenario (Figure 1C), if pair 1-5 resulted negative and pair 2-3 resulted in positive, electrode 1 would be labelled non-eloquent and electrode 2 would be labelled eloquent. Thus, if an electrode tested positive in at least two neighbouring pairs in any direction (i.e., horizontal, vertical or diagonal), the electrode was considered eloquent.

**Figure 1:**
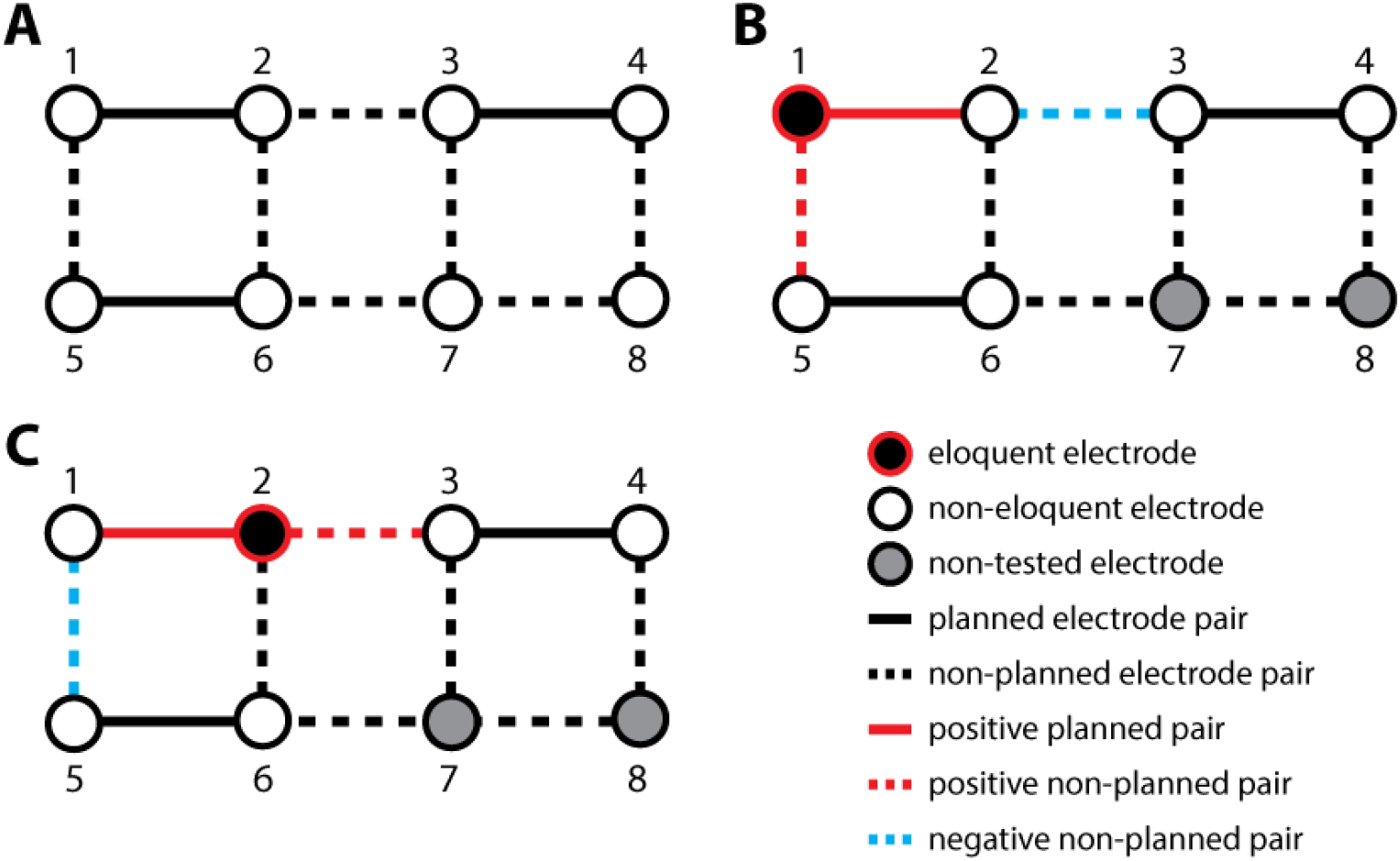
Schematic representation of two possible ESM cases in a toy example of a 2 by 4 electrodes ECoG grid. **A)** Planned stimulation pairs. **B)** Case one, in which electrode 1 is found to be eloquent after symptoms are produced during electrical stimulation at pair 1-2 and 1-5, but not 2-3. **C)** Case two in which electrode 2 is found to be eloquent after symptoms are produced at pair 1-2 and 2-3 but not 1-5.

The stimulation was performed with a constant current stimulator (Osiris cortical stimulator, Inomed, Emmendingen, Germany) with the following stimulation settings: trains of square wave pulses, pulse frequency 50 Hz, pulse duration 0.2 milliseconds, with a train duration of 3 to 7 seconds (depending on the function tested and the specific patient conditions). The current was increased in (1-)2 mA steps to a maximum of 15 mA (e.g., 1 mA, 3 mA etc.). During ESM for language testing, participants performed a reading task and, if reading was impaired, testing was extended with additional tasks (e.g., naming, counting, tongue movement task). ESM was stopped if any of the following end-points was reached: 1) a clear and reproducible generation of neurological symptoms, 2) impairment of any of the performed (cognitive) tasks, or 3) reaching the maximum stimulus intensity of 15 mA without causing symptoms, deficits or after-discharges. This procedure was then repeated at the next planned electrode pair. Stimulated electrodes that resulted in neurological signs or symptoms or cognitive impairment were labelled as *eloquent* and assigned to one of nine functional categories defined by the clinical neurophysiologists (motor, sensory, mixed sensorimotor, language temporobasal, language Broca, language Wernicke, emotion, visual, and auditory). This categorization was used as the ground truth against which our identification results were compared during the subsequent analyses.

All tests were performed in the epilepsy-monitoring unit (EMU) by an expert clinical neurophysiologist (V.H.K.M. or D.M.W.H). The selection of electrode pairs for stimulation and the total number of electrodes stimulated per session was decided individually for each participant by the clinical neurophysiologist. For the remainder of the text we use the term ‘eloquent’ interchangeably to refer to areas of cortex and to ESM positive electrodes, under the assumption that those electrodes correspond to eloquent cortex.

### 2.4. Delayed match-to-sample task

The stimuli and tasks were designed to test hypotheses about the processing of acoustic properties in the language network under varying conditions of attention, yet we reasoned that, considering the task properties, the same data might also be useful to test our current hypothesis.

A trial consisted of a speech sound stimulus comprising a spoken consonant-vowel syllable of 340 milliseconds duration followed, after a jittered interval of 550-750 milliseconds, by a written cue displayed for 500 milliseconds. After the presentation of the written cue, a 1,500 milliseconds period was allowed for a button response followed by an inter-trial interval (ITI) of 1,000 milliseconds as baseline before the next trial began (Figure 2 B). The written cue was either a complete syllable, a vowel, or consonant letter, depending on the specific block. The cue either matched or did not match the previous sound stimulus. Participants were asked to compare the syllable sound with the written cue and respond ‘match’ or ‘mismatch’ as quickly and accurately as they could. The stimuli matched in 50% of the trials, and ‘match’/‘non-match’ trials were balanced across conditions and randomized per participant. The identical sound stimuli were also presented in a passive listening condition in which sound onsets were jittered by 900-1100 ms while participants held their gaze steady on the computer screen. Stimuli were presented in the epilepsy-monitoring unit using a laptop computer with built-in open-field speakers. Stimulus and behavioural event triggers were sent to the clinical data recording equipment via a parallel port to the Ethernet interface box (Ethernet-102 V2, Braintronics B.V., Almere, Netherlands). All behavioural responses, stimuli, event identities, and timings were presented and logged using Presentation (Neurobehavioral Systems; www.neurobs.com, RRID: SCR_002521). The task was performed in 2 to 6 sessions per patient, in which trials were grouped into 4 blocks of 54 trials, with each block representing a different attention condition. Sessions lasted approximately 15 minutes and 1 to 2 sessions were performed per day. The total experiment time across sessions was between 30 and 90 mins per patient.

**Figure 2:**
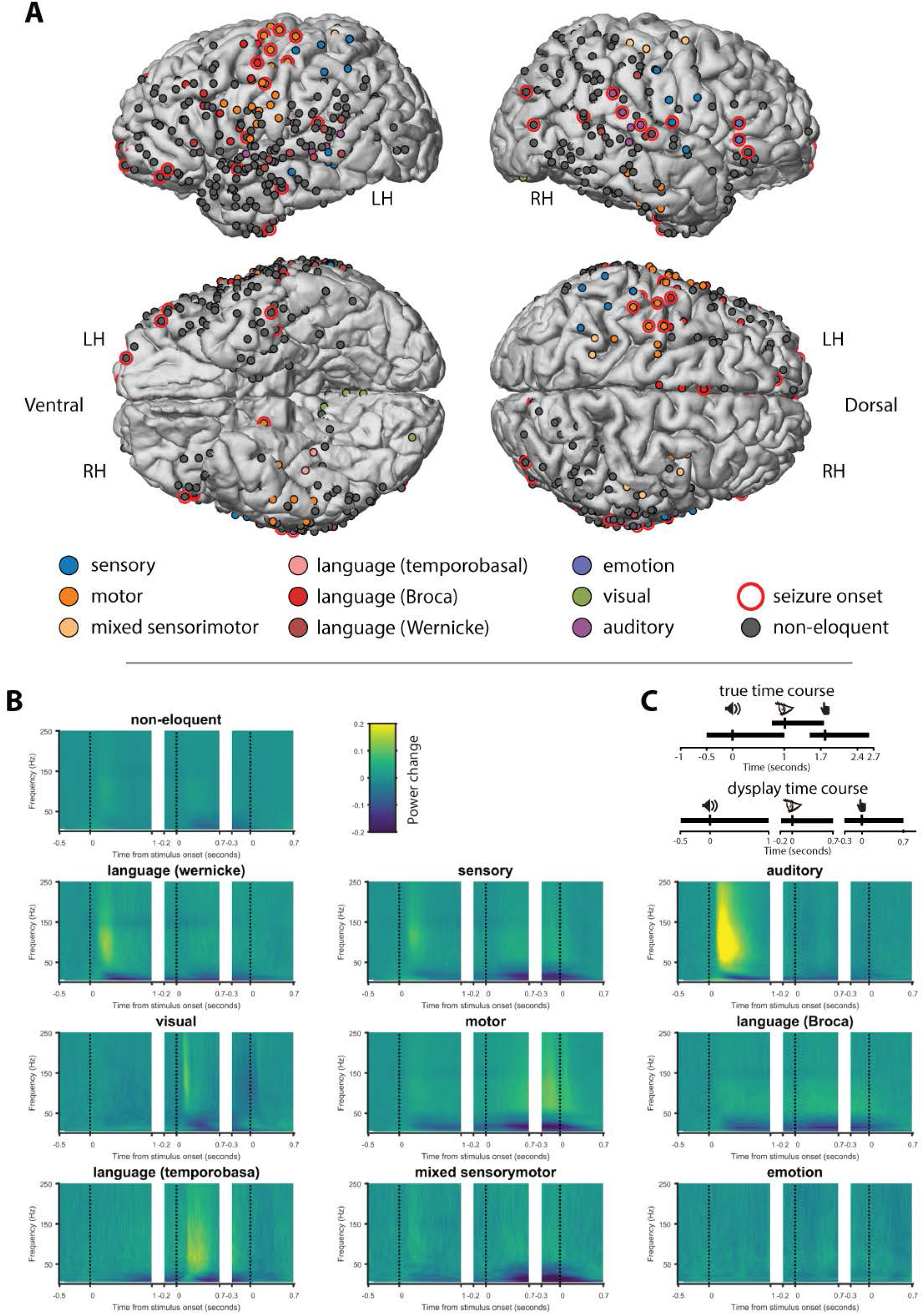
Electrical stimulation mapping and ECoG responses. **A)** Electrodes tested during electrical-cortical stimulation mapping (ESM) in the 10 patients projected in the common space. All electrodes are colour coded according with the ESM category or in black when no category was assigned. Brains represent the 3D mesh reconstruction of the grey matter-cerebrospinal fluid (CSF) boundary from the FreeSurfer average brain. LH: left hemisphere, RH: right hemisphere, Ventral: ventral (caudal) view of left and right hemispheres, Dorsal: dorsal (cranial) view of left and right hemispheres. **B)** Illustration of the time course in the raw trials (upper cartoon, where events are represented at their average time lag from sound onset, individual trials were jittered) and the ‘exaggerated’ time between the events used for display time (lower cartoon). **C)** Time frequency representations (TFR) of time periods of the delayed match-to-sample task from all tested electrodes grouped by the ESM category. The TFRs represent the three different trial events (i.e., sound onset, letter onset, and button press), which are illustrated by the three vertical dotted lines in each TFR).

### 2.5. ECoG data collection

ECoG signals were recorded at 2,048 Hz sampling rate using Brain RT software (version 2.0.3164.0, OSG BVBA, Rumst, Belgium). One or more 64-channel Brainbox EEG-1166 amplifiers (Braintronics B.V., Almere, Netherlands) were used to record from subdural electrodes (Ad-Tech Corporation, Racine, WI, USA) that consisted of platinum alloy discs embedded in a flexible silicon sheet. Electrodes had an exposed surface with a diameter of 2.3 mm. The electrodes were arranged in strips or grids with interelectrode center-to-center distance of 10 mm. As common reference, an inactive scalp electrode located over the forehead was used at the start of the recording and was in most cases changed to a relatively silent (i.e., not showing any epileptic activity during seizures) implanted electrode after the first seizure was recorded.

### 2.6. Data pre-processing

Data were analysed in MATLAB (R2016a version 9.0.0.341360; The Mathworks Inc.; Natick, MA, USA) using the FieldTrip toolbox (Oostenveld et al., 2011) and custom scripts. Data were first cut into epochs from 1 second before the sound onset until 1 second after the behavioural response with a maximum data length of 8 seconds. We then applied a discrete time filter at 50, 100, and 150 Hz to remove line noise and down-sampled the data from 2048 Hz to 500 Hz. Data were re-referenced to the average signal recorded in all electrodes, after excluding electrodes with high noise.

### 2.7. Time-frequency analysis

We calculated time-frequency representations of ECoG signal power in 2 Hz steps between 6 and 250 Hz using Hanning tapers of length equal to 7 times the cycle length and 10 ms step size. Power was expressed as the normalized change, where the change was defined as the difference between post-stimulus time window and pre-stimulus ‘baseline’ period −100 to −700 ms before sound onset. This baseline period was used to calculate power changes for all the events (sound, written cue and motor response) in the trial time window. The mean (not time resolved) spectral response was calculated from the normalized change from individual trial time frequency representations (TFRs), from the sound onset to 0.5 seconds after the behavioural response. To better represent the spectral response around each trial event, TFRs were aligned and cut around the onset of each event. Thus, we represent −0.5 to 1 second around the onset of the sound, −0.2 to 0.7 seconds around onset of the letters and −0.3 to 0.7 seconds around the onset of the behavioural response (Figure 2B).

### 2.8. Band-pass filtering and power calculation

For frequency band specific analysis we filtered at the classical alpha band (8 to 12 Hz) and beta band (15 and 25 Hz). We selected broadband gamma as 55 to 200 Hz, avoiding the possible line noise at 50 Hz. Power was calculated as the absolute value of the Hilbert transform. The resultant time courses were converted to normalized change in power from the baseline period (−0.7 to −0.1 seconds from sound onset). For time course analysis, individual trials were cut and aligned around each trial event, as described above for TFRs. For non-time resolved analysis we first calculated average power of individual trials from 0 to 0.5 seconds before taking the average across trials.

### 2.9. Generalized linear model (GLM) fitting and validation

Generalized linear models (GLM) are a generalization of linear regression models given by two main changes in the linear regression models, first the random component (Gaussian) is replaced by the exponential family of distributions and second, the link function (X^T^*β*) is replaced by a family functions (McCullagh and Nelder, 1998; Nelder and Wedderburn, 1972). This generalization process is achieved by an iterative weighting of linear regressions that are used to obtain maximum likelihood estimates of the parameters. Given that the power modulation response might contain nonlinearities, the use of a GLM allows identifying the optimal weighting for each frequency band combination, thus increasing the performance in eloquent-electrode’s identification.

We used a GLM with binomial likelihood and logit link function (McCullagh and Nelder, 1998), where for each electrode the dependent variable (y) was the ESM response (eloquent and non-eloquent) and the predictors were the ECoG power change in each frequency band. Once the model was estimated, we could determine the optimal weighting of the power change at each frequency band as regression coefficients to predict the ESM response. Since calculations of AUC using the same data used for GLM estimation would result in an inflation of type-I errors, we estimated the GLM coefficients on a subset of electrodes and determined the area under the receiving operating characteristic curve on the remaining portion. Thus, we first partitioned the eloquent and non-eloquent electrodes categories into k disjoint datasets and used, the data contained in k-1 partitions (i.e., leaving one partition out for testing later the mode’s prediction) to estimate a GLM having ECoG power change in each frequency band as independent variables x, and eloquent or non-eloquent ESM categories as the dependent variable y, (y being 0 for non-eloquent electrodes, and 1 for eloquent electrodes). Both quadratic and interaction terms were included in the estimation in order to benefit from possible non-linearities in the ECoG response (power change) and for possible interactions among the two. The model was then used to predict the ESM response in the left out partition. The whole procedure was repeated for all the partitions (i.e., k-fold cross-validation, in our case we used k = 10). The resulting prediction, concatenated across all the test partitions, was compared with the ground truth ESM by means of area under the receiver operator characteristic (ROC) curve. Finally, the whole procedure was repeated 20 times randomly selecting the partitions and the results were averaged. When considering each frequency band alone, we performed a similar analysis including the quadratic terms, but using only the one predictor of interest.

### 2.10. Area under the receiver operator characteristic curve analysis

The area under (AU) the receiver operator characteristic (ROC) curve or AUROC is a measure of discriminative ability for continuous predictors such as prediction models. The ROC curve is drawn in a space that represents sensitivity (true positive divided by the sum of true positive and false negative) against 1-specificity (true negative divided by the sum of true negative and false positive). The curve connects the combinations of sensitivity and specificity for all possible risk thresholds. AUC is the magnitude of the area under this curve. The models being evaluated are the ones generated by the GLM for the different frequency bands and their combination. The threshold of AUC needed to determine the usability of a diagnostic test depends on the planned use of the prediction model and the cost or morbidity generated by the proposed intervention. In the present study, the proposed intervention consists of an inexpensive additional cognitive task in combination with an automatic computer analysis of the cortical frequency power changes. Thus, the required threshold for the AUROC is relatively lower compared to more expensive tests such for example ESM (Kundu et al., 2016).

We aimed to calculate the diagnostic ability of the three frequency bands, and of their combination. We did not implement a diagnostic test, for which we believe more data, and an optimized behavioural task, would be required. Previous studies have quantified the sensitivity and specificity of changes in cortical power. However these values are determined by both the diagnostic ability and the discrimination threshold applied. We calculated the area under the curve (AUC) from the receiver operating characteristic (ROC) curve (Green and Swets, 2000). This procedure calculates the ratio of sensitivity to specificity for all potential discrimination thresholds. The resulting AUROC represents the maximum performance (i.e., correct discrimination) of an ‘ideal observer’ using the optimal threshold. The diagnostic ability may be considered significant if the AUROC is different from chance (50% correct). In our analysis, AUROC was higher than 0.5 when the response (i.e., power change within a given frequency band) in eloquent electrodes was greater than in non-eloquent electrodes, and less than 0.5 when the response in eloquent electrodes was lower than the response in non-eloquent electrodes. Significance was therefore tested using a two-tailed test, where the null distribution was empirically calculated by randomly permuting (1000 times) the labels between eloquent and non-eloquent electrodes and calculating the corresponding AUROC from the permuted data.

### 2.11. Bootstrap procedure for statistical testing receiver operator characteristic

Statistical differences among the different models resulting from the GLM fitting (each frequency band and their combination-see previous section) were tested using a bootstrap analysis. We constructed a distribution of AUROC values for each model by creating multiple datasets of the same size as the original by randomly drawing electrodes with replacement. On each dataset we computed the AUROC as in the previous step for each model and obtained a distribution of values across 1000 bootstrap samples. Statistical comparisons were then carried out by comparing the differences between models. We considered the difference between two models as significant when one model outperformed the other in more than 95% of the cases.

### 2.12. Function-specific analysis

We were interested to study whether different functional areas might exhibit characteristic power spectra, however we were limited by the low numbers of electrodes representing some eloquent ESM categories. We therefore grouped electrodes into two loosely defined categories which we call ‘receptive’ and ‘expressive’. Those electrodes labelled as *auditory, visual, sensory, language-Wernicke and language-temporobasal* were grouped into the ‘receptive’ category as being more functionally involved in processing incoming sensory events. Those labelled as *motor, mixed-sensorimotor and language-Broca* were grouped into the ‘expressive’ category as being more involved in generating behavioural responses. Two neighbouring electrodes in one patient labelled as ‘*emotion’* were excluded from the function-specific analysis as they did not seem to fit in either category.

### 2.13. Attention dependent analysis

To test the influence of attention, we compared power changes in the different frequency bands of interest (broadband gamma, alpha and beta band) from the passive listening condition to power changes from the different frequency bands from the DMTS task. To match the number of trials performed in each task we used data from only one DMTS condition, where the written cue represented full syllables, and used a comparable time window in both tasks (−0.5 seconds to 0.9 seconds from the sound onset).

### 2.14. Probabilistic map of eloquence

We considered how our analysis could to be clinically useful without providing a diagnostic test. A probabilistic map that shows the likelihood of eloquence for each electrode obtained before the ESM could be used to optimize the planning of the sequence of electrode pairs for ESM testing. We took GLM calculated probabilities for each patient separately using only data from the remaining patients to calculate the GLM beta weights. This procedure represents a special case of our *standard* analysis in which data was separated into training and test datasets: In the *standard* analysis the separation was done randomly, thus data from different patients could be included for training and the same could be done for testing; in the *current* analysis the data separation was performed according to the patient identity. In this way we show how a probabilistic map of likelihood of eloquence in ESM can be generated for an individual patient without reference to the ESM results of that patient. This procedure allows, in a new patient, the creation of a probabilistic map of eloquence using only data from the DMTS task that is acquired before the ESM, enabling the incorporation of the probabilistic map in the planning of the ESM.

## 3. Results

### 3.1. Relationship between electrical cortical stimulation and frequency modulation

As a descriptive analysis, we selected all the electrodes from all ESM categories and projected them into the common brain space from FreeSurfer to see the distribution and location of the electrodes from the different patients (Figure 2A). We registered in total 129 eloquent and 460 non-eloquent electrodes (Table 1). Eloquent electrodes were located bilaterally over the temporal neocortex (superior and middle temporal gyri), over the inferior frontal gyrus, and over the pre- and post-central gyri.

**Table 1:** Electrode details. Number of electrodes in each ESM category, total number of eloquent, seizure onset, and non-eloquent electrodes per patient.

| Ptn | all elec | non-eloquent | total eloquent | seizure onset | sensory | motor | sensory-motor | language (wernicke) | language (Broca) | language (temporobasal) | auditory | visual | emotion |
| --- | --- | --- | --- | --- | --- | --- | --- | --- | --- | --- | --- | --- | --- |
| 1 | 40 | 36 | 4 | 5 | 0 | 0 | 2 | 0 | 0 | 1 | 1 | 0 | 0 |
| 2 | 39 | 39 | 0 | 2 | 0 | 0 | 0 | 0 | 0 | 0 | 0 | 0 | 0 |
| 3 | 40 | 29 | 11 | 0 | 4 | 0 | 0 | 1 | 0 | 0 | 6 | 0 | 0 |
| 4 | 74 | 59 | 15 | 12 | 0 | 8 | 0 | 0 | 7 | 0 | 0 | 0 | 0 |
| 5 | 31 | 31 | 0 | 0 | 0 | 0 | 0 | 0 | 0 | 0 | 0 | 0 | 0 |
| 6 | 41 | 27 | 14 | 8 | 4 | 2 | 0 | 1 | 0 | 2 | 3 | 0 | 2 |
| 7 | 79 | 43 | 36 | 7 | 5 | 14 | 2 | 7 | 7 | 0 | 1 | 0 | 0 |
| 8 | 74 | 63 | 11 | 3 | 1 | 0 | 3 | 1 | 0 | 0 | 2 | 4 | 0 |
| 9 | 92 | 69 | 23 | 7 | 2 | 14 | 3 | 3 | 1 | 0 | 0 | 0 | 0 |
| 10 | 79 | 64 | 15 | 3 | 4 | 8 | 0 | 0 | 0 | 0 | 0 | 3 | 0 |
| **Total** | **589** | **460** | **129** | **47** | **20** | **46** | **10** | **13** | **15** | **3** | **13** | **7** | **2** |

We represented the change in power spectral density, relative to the baseline period (−0.7 to −0.1 seconds from sound onset) across time (time-frequency representation, TFR) of the ECoG signal during each epoch of the DMTS task (Figure 2B). We observed that ESM categories exhibited characteristic frequency response patterns during the task. For example, electrodes labelled during ESM as *‘auditory’* exhibited the strongest activity increase in gamma and decrease in low frequency activity (alpha and beta power, LFA) after sound onset. Likewise, electrodes labelled as ‘*visual’* exhibited an increase in gamma and a decrease in LFA after letter onset. Electrodes labelled as ‘*motor’* exhibited an increase gamma activity and a decrease in LFA around the button press. Interestingly, ‘*language temporobasal’* and ‘*language Broca’* electrodes seemed to be active after the sound onset and after letter presentation possibly pointing towards covert rehearsal of the sound and silent reading of the letters. For further analysis, we grouped all electrodes labelled as eloquent for comparison with all electrodes that were tested using ESM but labelled as non-eloquent.

### 3.2. Receiver Operating Characteristic (ROC) curve analysis

We compared the power spectrum of eloquent and non-eloquent electrodes’ ECoG during the DMTS task. We observed that power was generally increased relative to baseline for frequencies above 50 Hz, and generally decreased for frequencies below 30 Hz (Figure 3A). This effect was larger in eloquent electrodes (red) than non-eloquent electrodes (blue). To test the diagnostic ability of this difference we calculated AUROC (Figure 3B) and applied a permutation test for statistical assessment. AUROC was significant (thicker line, alpha = 0.05, double sided test, uncorrected for multiple comparisons) for frequencies between 50 Hz and 180 Hz and for frequencies between 6 and 30 Hz, indicating significant diagnostic ability, with a performance of 65 to 70% correct for an ideal observer.

**Figure 3:**
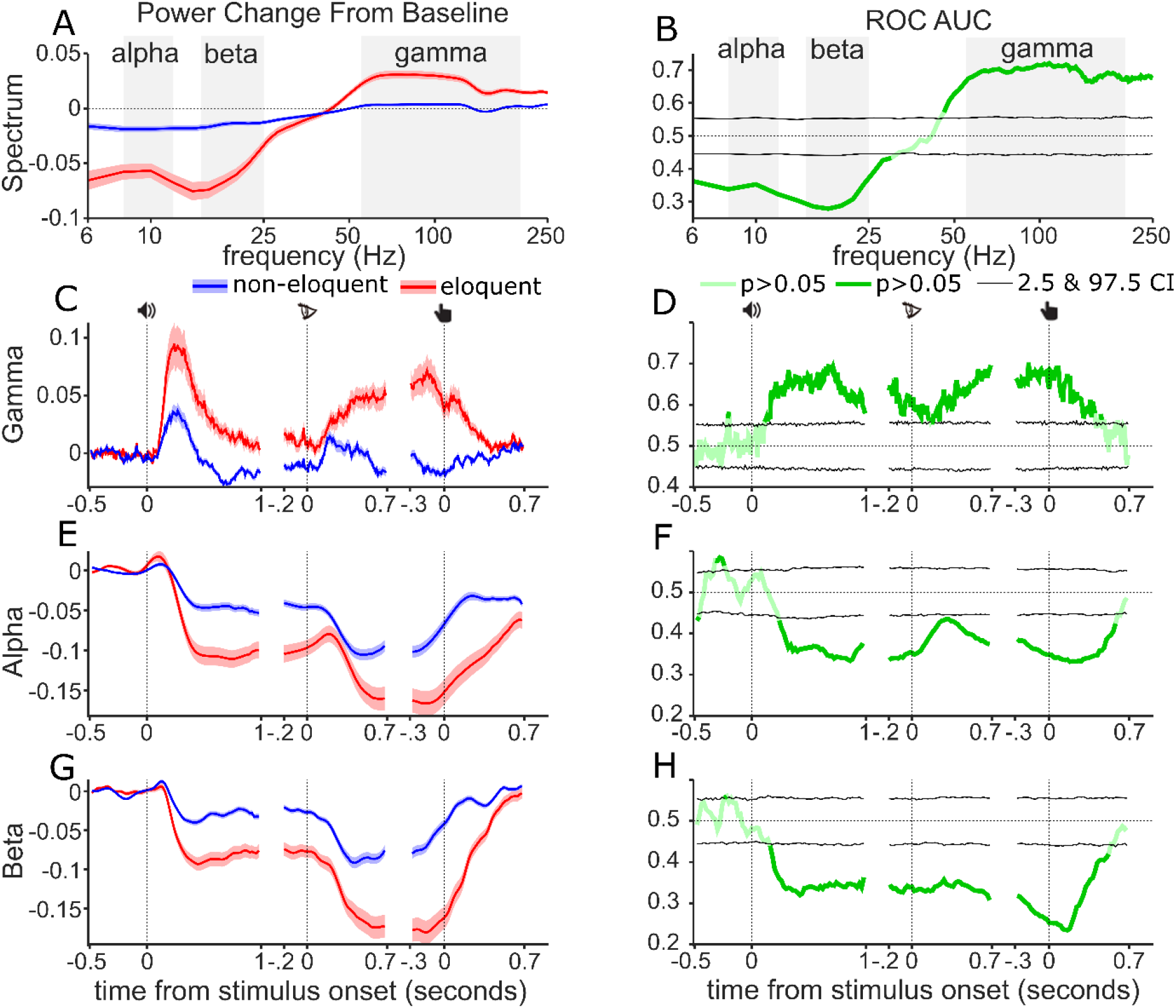
AUROC analysis. **A)** Normalized change in ECoG signal power spectral density from baseline from eloquent (red) and non-eloquent electrodes (blue). Shading width shows standard error, centre line shows mean. **B)** AUROC values across frequencies, pale green, non-significant, dark green significant values. Black lines show 0.25 and 97.5 percentiles from permutation distribution. Dotted line shows chance performance. Background shading indicate filter boundaries for alpha, beta and gamma. **C)** Gamma power modulation during the three events of the delayed match-to-sample task (icons indicate sound onset, letter onset, and button press). Solid lines show mean, shadow width; standard error. **D)** AUROC values across time for gamma. **E&F)** Alpha power and AUROC values over time, as in C&D. **G&H)** Beta power and AUROC values over time, as in C&D.

To represent the time courses of gamma- alpha- and beta bands over the trial we filtered the ECoG signal in the respective bands (grey background shading, Figure 3A&B) and calculated the absolute value of the Hilbert transform, which was subsequently represented as the normalized change against pre-sound baseline period. We found stronger gamma power increase in eloquent as compared with non-eloquent electrodes (Figure 3C) during the three different DMTS task epochs. The AUROC values across time (Figure 3D) were larger than 0.5 and significant for all three events in the DMTS task.

Alpha power (Figure 3E) showed a decrease for all the three DMTS task events with an overall pattern for both eloquent and non-eloquent electrodes (Figure 3E) and a stronger decrease from baseline in eloquent electrodes compared with in non-eloquent electrodes. The AUROC values were significantly below chance for all three events (Figure 3F). A similar pattern was observed for beta power, (Figure 3 G&H) however, the decrease in power was especially prominent around the button press event. AUROC values were generally lower, indicating greater diagnostic ability, than for the alpha band, especially around the button press event.

Next we calculated AUROC values across time using the weights from the GLM, including 10-fold cross-validation (Figure 4A). For smoothing, and to reduce processing time, we used a sliding window using a 100ms window and 50ms step size. The GLM based AUROC values for gamma-only closely matched standard AUC values (Figure 4A, green line). For alpha-only (red) and beta-only (blue) the GLM based AUC values also closely matched the standard AUROC, except that values were above 0.5 rather than below due to the fitting procedure, see section 2.10. Interestingly, time courses for the three individual band models AUROC values peaked at different time periods of the trial. The gamma-only model performed best during the sound epoch while the beta-only model performed best during the button press epoch. The alpha-only model performed generally worse than the other models, but interestingly outperformed the gamma-only model around the letter onset and late after the button press. The combined GLM AUROC values (Figure 3G–dashed lines) tended to be either higher than, or equal to whichever individual curve was highest at any given time point. This was especially true for the gamma&beta model (blue dashed lines), although the alpha&gamma (red dashed) model also tended to outperform the gamma-only model. Interestingly the three-band model (black dashed) did not outperform the gamma&beta model.

**Figure 4:**
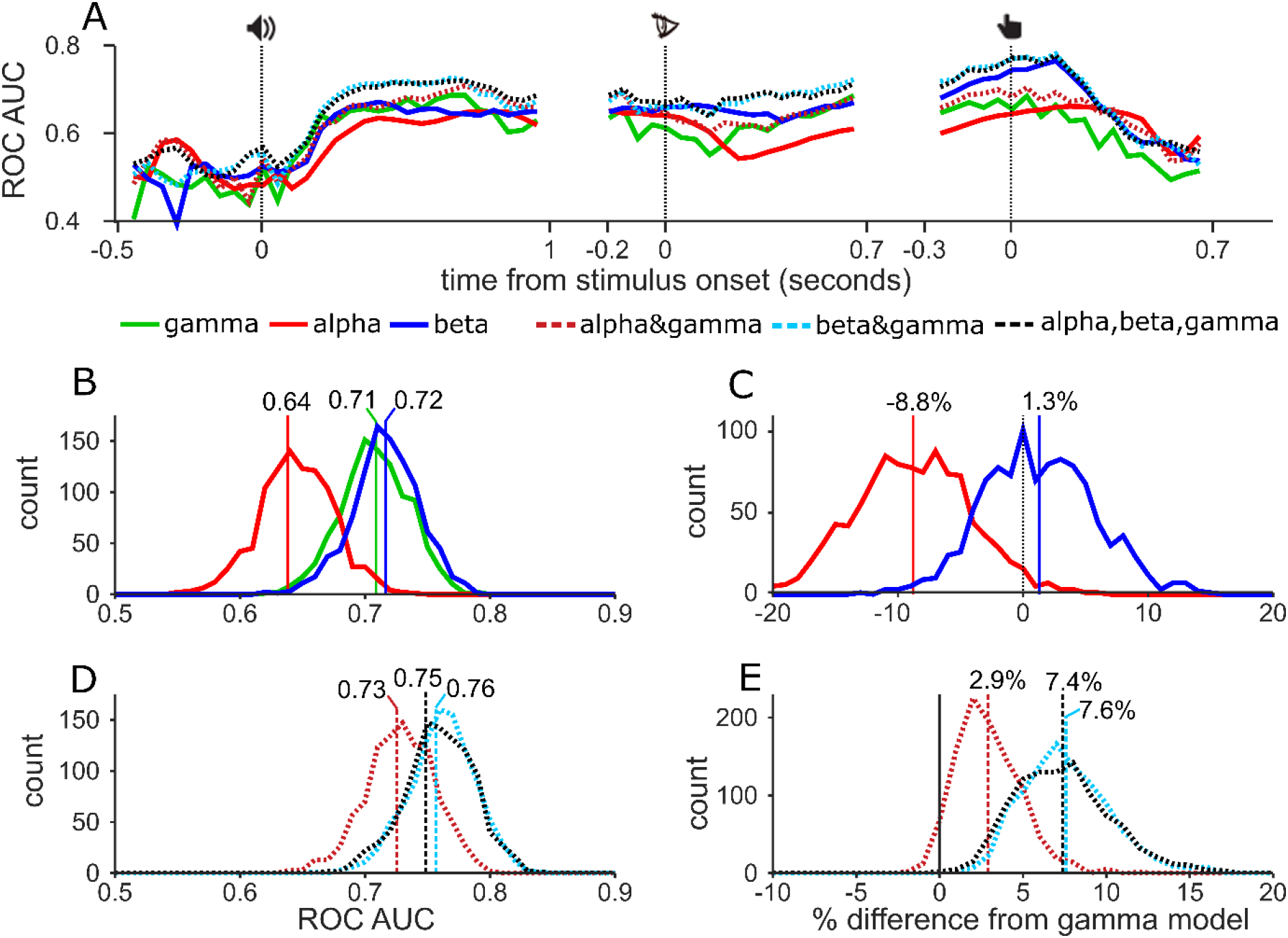
**A)** AUROC values using GLM approach across time for alpha band, gamma Beta (solid lines), and the combinations (dashed lines). **B)** Distribution of the AUROC values from 1000 bootstrap samples for single models. Vertical lines show ROC AUC values from all electrodes without resampling. **C)** Pairwise comparison from bootstrap distribution of alpha (red) and beta (blue) AUROC values with gamma values. Vertical lines show mean percentage change from gamma performance. **D&E)** Distribution of AUROC values for combination models, as in B. **E**) pairwise comparisons of combination models with gamma-only model.

To statistically test the AUROC results of each model against chance performance and against each other, we first reduced the dimensionality of the data by calculating average power for each electrode, collapsing the data over the time dimension. Using this data we conducted a bootstrap analysis in which we constructed a distribution of 1000 AUROC values for each model by creating multiple datasets of the same size as the original by randomly drawing electrodes with replacement. For each dataset, and the original, we compute the AUROC for each model using 10-fold cross-validation. Histograms in Figure 4B show the distribution of AUROC values of the single band models, vertical lines show the AUROC values from the original dataset. All three models performed better than chance, with 100 percent of the resampled datasets returning AUROC values greater than 0.5. Of the three models the alpha-only model (red) performed the worse, with an AUROC of 0.64 in the original dataset, the gamma only (green) and beta only (blue) performed similarly and the distributions substantially overlapped. The gamma-only approach has previously been used most widely, therefore to test the value of alpha- and beta bands we made pairwise comparisons of those models against the gamma-only model. Figure 4C. The alpha-only model was worse than the gamma-only model in 98% of datasets, which we consider to show a significant difference. The mean performance reduction was 8.8% (vertical line). While the beta-only model offered a slight performance advantage on average (1.3%) this was not significant. We next compared the performance of the combination based models (Figure 4D). The three combination models generally performed better than the individual-band models although the alpha&gamma (red dashed) model performed the worst. The beta&gamma (blue dashed) and three-band (black dashed) model performed approximately equally. In pairwise comparisons with the gamma-only model, all three models offered enhanced performance although this just failed to meet the threshold of significance for the alpha&gamma model (94% of datasets improved). The beta&gamma and three band models both performed significantly better than the gamma-only model (in 100% of datasets) offering respectively an average 7.6% and 7.4% improvement. Interestingly, their distributions completely overlapped indicating that there was no advantage to including the alpha band to the beta&gamma model.

### 3.3. AUROC performance depends on functional category

Alpha-, beta- and gamma band activity have been ascribed different functional roles, with gamma associated with feedforward processes and alpha associated with feedback processes (Scheeringa and Fries, 2017). The beta band has been associated with motor activity, whereby beta power drops in preparation for a motor response. Thus, different frequency bands may have relatively different importance in different cortical areas, depending on the area’s place in the cortical hierarchy (Felleman and Van Essen, 1991). Specifically, we can anticipate that receptive areas, being early in the hierarchy, may have a high dependence on gamma, while expressive areas, being late in the hierarchy may be more dependent on alpha and beta. Importantly to our current question, this implies that patients with many expressive electrodes will be poorly mapped using gamma. For those patients, alpha or beta may be more informative, or their combination with gamma may give a greater improvement. To test this hypothesis we divided all eloquent electrodes into two loose categories which we defined as *expressive* and *receptive* and repeated our analysis (Figure 5) for each group separately. Notice that this grouping was intended as a procedural means to split the data into denominated groups. With the possible exception of primary areas, all cortical areas have roles in both processing stimuli and generating responses, thus the distinction between *expressive* and *receptive* areas is only an approximation.

**Figure 5:**
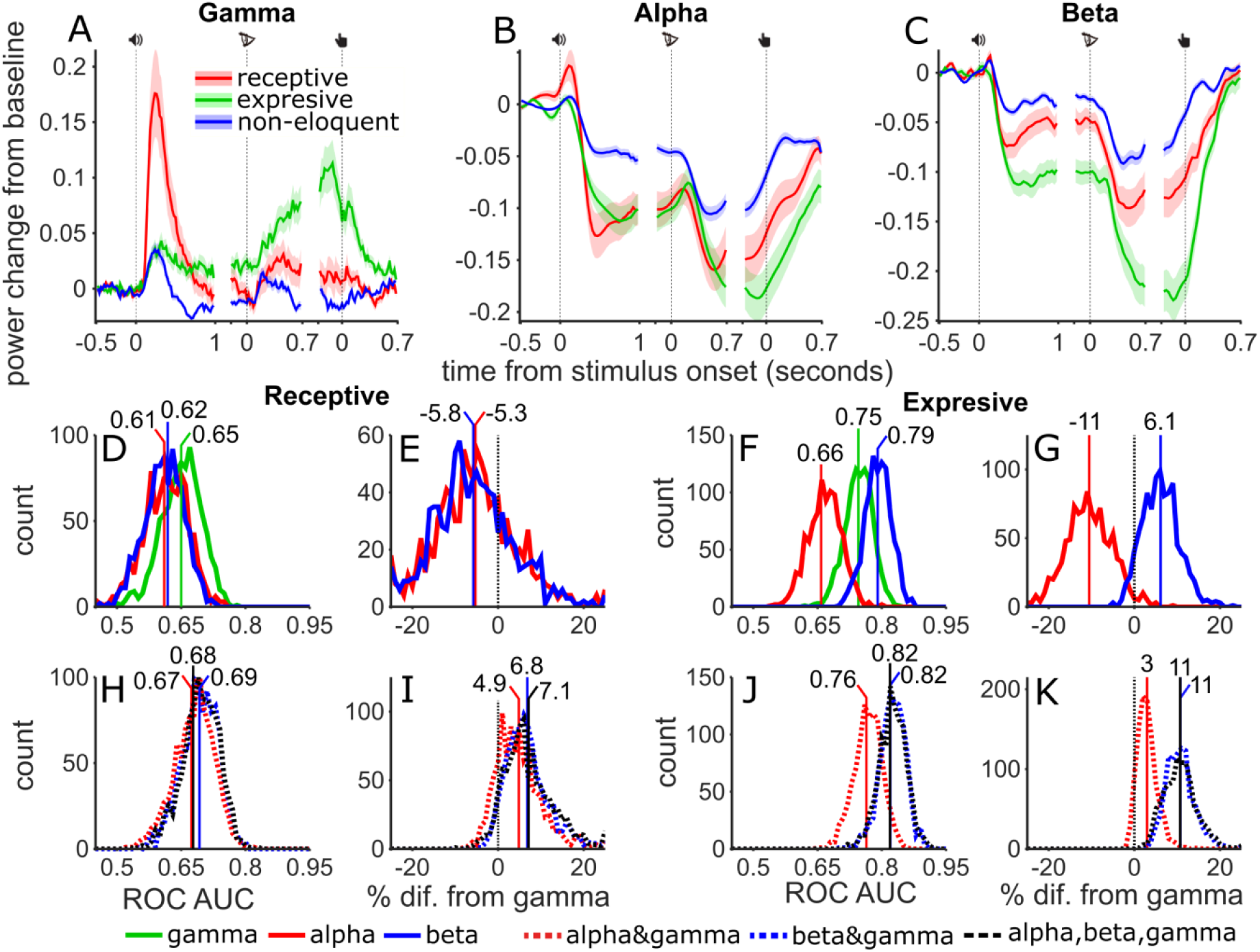
AUROC analysis per functional category. **A,B,C)** Gamma, Alpha and Beta power modulation for the three subtypes of electrodes (receptive, expressive, and non-eloquent) during the three events of the delay match-to-sample task. **D)** Distribution of the AUROC values from 1000 bootstrap samples for single models (solid lines) and receptive channels. **E)** Pairwise comparison from bootstrap distribution of alpha (red) and beta (blue) AUROC values with gamma values for receptive channels. H&I) **F & G)** Same as D & E for expressive channels. H to K, same as D to G using combined models (dashed lines).

Descriptively, gamma (Figure 5A) after sound onset showed power increase from baseline for all electrodes that peaked at ∼100 ms after sound onset. Receptive electrodes (red line, line shading shows standard error) showed the highest power followed by expressive electrodes (green). Non-eloquent electrodes (blue) showed the weakest response. After letter onset, gamma showed an increase in power for all electrodes. While power in receptive electrodes peaked soon after letter onset, power in expressive electrodes continued to increase reaching a peak shortly before button press. Non-eloquent electrodes showed low power after a weak response to letter onset and button-press. Alpha power (Figure 5B) was suppressed in all channels with greater suppression for eloquent than non-eloquent channels. Interestingly, alpha power suppression was approximately even in both expressive and receptive channels for the majority of the time course, with only a slight difference as the end of the trial where expressive channels showed somewhat greater suppression. By contrast, beta band (Figure 5C) suppression was considerably stronger in expressive channels than receptive channels at all time points of the trial after sound onset. With our stated caveat about the distinction between expressive and receptive areas, these data supported our grouping as ‘*receptive*’ electrodes seemed most involved at the start of the trial during stimulus processing, and ‘*expressive*’ electrodes seemed most involved at the end of the trial near the behavioural response.

We calculated AUROC values for the three bands separately and for the three combination models using the time-averaged responses and tested significance of the difference using the bootstrap method. In *receptive* (Figure 5D) electrodes the distribution of gamma (green) AUROC values was higher than the alpha-only (red) or beta-only (blue) models, however the distributions heavily overlapped. Pairwise comparisons (Figure 5E) showed that this difference was not significant (alpha- and beta-only models overlapped the performance of gamma-only). In *expressive* electrodes (Figure 5F) the beta-only model outperformed both gamma-only and alpha-only models. Pairwise comparisons (Figure 5G) showed that this difference corresponded to a 6.1% increase but did not pass significance (93% of datasets). The alpha-only model performed significantly worse than the gamma-only model, corresponding to 11% drop.

Analysis of the combination models showed that in *receptive* channels all three models offered approximately equal performance (Figure 5H), which was somewhat better than the individual band models, however pairwise comparisons (Figure 5I) showed that this improvement failed to meet the threshold for significance. Among *expressive* channels the combination models offered better performance (Figure 5J) and a greater improvement over the gamma-only model (Figure 5K). In the alpha-gamma model mean improvement was 3% but failed to reach significance, while the beta-gamma and three band models both showed an improvement in 100% of dataset with a mean improvement of 11%. These results were in line with our expectation that *expressive* electrodes, being higher in the cortical hierarchy, would be most sensitive to low frequency power modulations. Regardless of the interpretation, taken together these results suggest that eloquent areas may be best mapped using gamma or beta-band power depending on the distribution of electrodes in an individual patient. However, irrespective of the ESM functional category, the combination of beta and gamma bands was reliably the best measure for the identification of eloquent electrodes.

### 3.4. The influence of attention on AUROC values

Most patients performed well in the task, however one patient performed poorly and withdrew from the experiment after only two sessions (out of a planned 5). This experience prompted us to question how well ECoG mapping would perform in a less demanding task, which might be particularly relevant when applying ECoG mapping in paediatric populations, patients with low cognitive ability or patients who are otherwise unable or unwilling to engage in cognitive testing. Our cognitive experiment included a passive listening condition in which auditory syllables were presented without any explicit task. We repeated our analysis using the ECoG response, from eloquent (red lines) and non-eloquent electrodes (blue lines), during the sound presentation in the active (solid lines, delayed match-to-sample task) tasks and passive condition (dashed lines, Figure 6). Gamma band (Figure 6A) responses were weaker in the passive task compared to active task for both eloquent and non-eloquent electrodes. Similarly, alpha and beta band suppression Figure 6B & C) was weaker in the passive task compared to the active task, especially late in the response.

**Figure 6.**
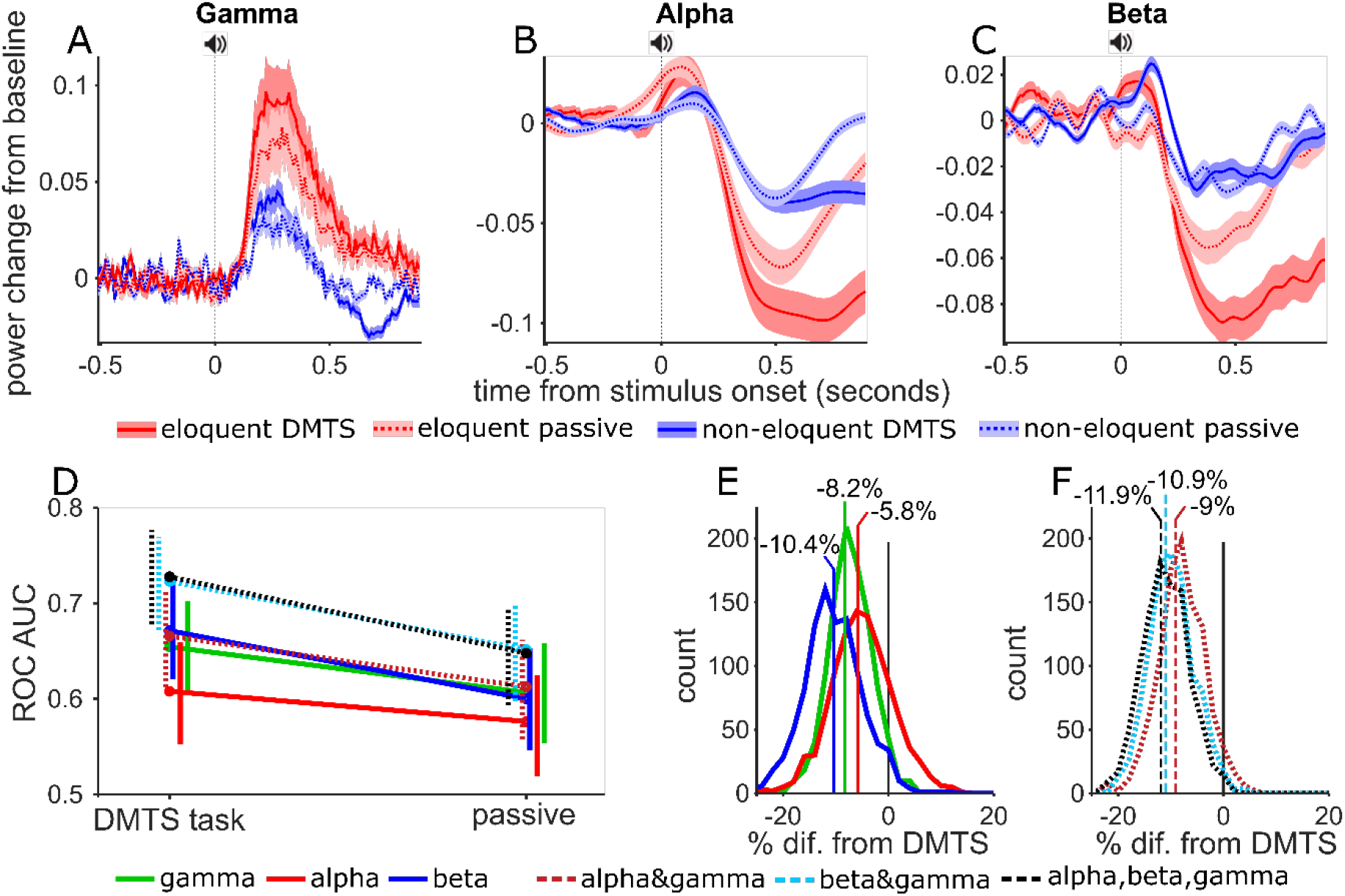
Attention effect on AUOC values. **A,B,C)** Gamma, Alpha and Beta power modulation across time for eloquent (red) and non-eloquent (blue) electrodes during the active (DMTS, solid lines, darker shading) and the passive (dashed lines, lighter shading) listening tasks. **D)** Effect of attention of diagnostic ability per model. Vertical lines show 5 and 95 percentiles from 1000 bootstrap samples, horizontal offset to aid visibility. E) AUROC values for all models, separately for active (DMTS) and passive task. **E)** Pairwise comparison between active and passive task per single band model. Negative values imply a better performance in the active task. **F)** Same as E but using combination models.

As before we calculated AUROC values of the GLM models and tested significance using the time-averaged responses, however here we used only the period from sound onset to 900 ms which could be compared for both tasks. In the DMST data AUROC values were unsurprisingly lower than in previous analysis (Figure 4H) where we had used the response during the entire DMTS task. However, when comparing the performance of the three bands and the combination, the pattern matched our previous findings. Comparing the two tasks (Figure 6D) showed that withdrawing attention from the stimuli in the passive task suppressed the performance of all models. In pairwise comparisons between the two tasks we found that performance of the models dropped by as much as 11.9% in the three-band models. The drop was significant (*p* < 0.05) for all models expect for the alpha-only model (*p* = 0.19).

These data showed that an active task which engages attention is required for optimal mapping using ECoG, with important implications for studies which attempt to classify eloquent cortex without explicit cognitive tasks (Vansteensel et al., 2013). Nevertheless, the GLM approach using gamma and beta band activity offered a significant improvement over single-band models. Notice, that this analysis was possible because we had collected passive data in all patients.

### 3.5. Probabilistic map of likely eloquence

We aimed to use the GLM fitting to calculate a map showing the probability that each electrode would be eloquent given the ECoG power modulation during the DMTS (i.e., GLM-ECoG). Such a map could be used to optimally plan the sequence of ESM mapping. For this analysis we used data from the full trial and full dataset (i.e. data presented in Figure 2H & I) and used the gamma&beta combination model. As in the main analysis we estimated GLM weights on a training dataset and applied those weights to a test dataset here training and testing was performed in splits containing data from all patients. However unlike in the main analysis, for the creation of the probabilistic map the training dataset comprised of all electrodes from all patients minus one and the testing dataset comprised of all electrodes from the remaining patient. Thus, this analysis could be performed for a new patient using data available before the ESM (i.e., data from the DMTS task). After calculating the prediction of all electrodes in the dataset the AUROC for this procedure was 0.73, as compared to 0.76 using the K-fold approach, in line with the results from the standard analysis.

The output of the GLM (i.e., GLM response) corresponds to a prediction of the binomial probability that an electrode is eloquent or not. To visualize the relationship between GLM prediction and the empirical probability of eloquence we binned electrodes into 10 equally spaced bins (with 50% overlap) according to the output of the GLM. Figure 7A shows the number of electrodes per bin (black line, rightward Y axis) and the proportion of those electrodes with a positive ESM response (blue line and coloured dots, leftward Y axis). The proportion of eloquent channels across the whole population is shown by horizontal line. We then showed how the GLM estimate could be used to map five representative patients. The probability of eloquence was represented by colour-coding the electrode locations, with darker-red colours indicating higher probability of eloquence (see dot colours in Figure 7A). For comparison, electrodes that were eventually labelled as eloquent by the ESM were marked with a central black dot while electrodes eventually labelled as non-eloquent were marked with a central white dot. For completeness, electrodes labelled as seizure onset were labelled with a red ring. Patient 4 and 8 appear to have a good match between ESM and ECoG prediction in that darkest coloured dots also have black centres. Eloquent areas were far from seizure onset zones allowing a safe resection. Patient 7 also had good match between ESM and the ECoG prediction however the overlap between eloquent areas and seizure onset zone precluded a safe resection. Patient 3 showed a good match between ESM and ECoG however notice that the estimated probabilities were generally low in this patient such that ESM positive electrodes were found at a level of ECoG response that would be negative in other patients. This observation indicates that a fixed diagnostic threshold applied to all patients may not be the ideal approach. Patient 2 showed the reverse pattern, in this patient no eloquent electrodes were found with ESM despite quite high ECoG responses. In this patient the seizure onset zone was located at the temporal pole, while ESM results do not contra indicate a full resection of the temporal lobe, the ECoG power modulation response seems to suggest active cortex above the 5th row of the grid. These results could be taken into account as additional source of information when planning a resection for the removal of epileptogenic tissue while preserving regions with high ECoG power modulation responses.

**Figure 7.**
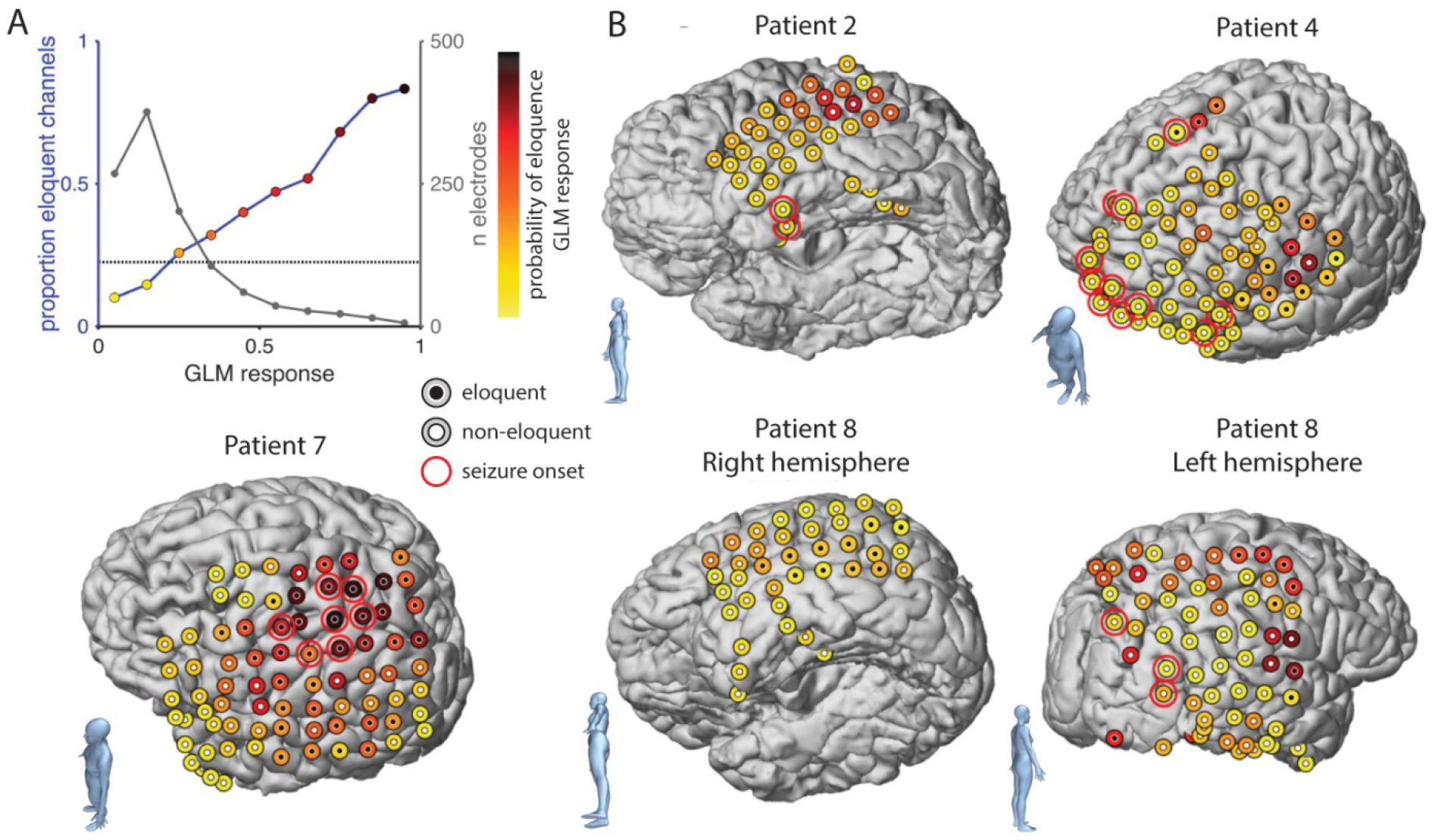
Probabilistic maps of likely eloquence. **A)** Relationship between GLM response (i. e., prediction of the binomial probability that an electrode is eloquent or not) and probability of eloquence (leftward axis) and numbers of electrodes (rightward axis). Dot colours indicate colour-scale used in other panels. **B)** Electrode locations projected onto the individual patient MRI in five example patients. Dot colour indicates GLM response, dot centres indicate ESM results, white: non-eloquent, black: eloquent, coloured: not mapped. Red rings indicate seizure onset zones. Blue bodies in the bottom left of each brain reconstruction indicate the brain orientation and observer view.

## 4. Discussion

We found that activity in the alpha, beta and broadband gamma frequency ranges could be used to identify eloquent cortex at above chance level, and with rates in line with previous reports (Arya et al., 2018). Combining frequency bands via a generalized linear model (GLM) enhanced the information given by each band alone, thereby increasing the prediction of eloquence. Combining the beta band with the broadband gamma was found to be more useful than combining the alpha band with broadband gamma. Time-resolved analysis showed that the different frequency bands alternated as the ‘better’ measure throughout the trial. Likewise segregating the signal from the eloquent electrodes into *receptive* and *expressive* groups showed that the relative performance of alpha, beta and broadband gamma depended on the cortical area, whereby broadband gamma gave better results in *receptive* areas and beta band gave better results in *expressive* areas in line with previous results reported by Wu et al. (2010). These analyses reveal that whether relatively lower frequency alpha/beta or higher frequency broadband gamma power are the ‘better’ measure, will depend on several factors, including the patient’s implantation scheme, the task performed, and the selected analysis window. Our GLM approach cuts through this complexity because, regardless of which band provided the better diagnostic ability, the combined approach allowed a performance at least as good as the best individual frequency band and typically offered an improvement.

Previous studies have investigated whether ECoG recording could be used as an alternative to ESM by testing whether a significant response could be used as a diagnostic criterion for identifying eloquent cortex. Here we investigated the diagnostic ability without implementing a diagnostic test. After demonstrating that ECoG responses can reliably predict the probability of an eloquent ESM response, we showed how a probabilistic map could be constructed that could be used to guide ESM. Guided ESM may reduce the time and effort from both the patient and the clinical team to stimulate all possible pairs of electrodes from an implanted grid. The time-consuming aspect of ESM becomes especially problematic as the number of electrodes increases, which is the case with the use of high-density ECoG grids. High-density grids have become more frequently used as they increase the spatial resolution of the mapping procedure, thereby increasing surgical precision and decreasing the risk of postsurgical neurological deficits (Escabí et al., 2014). However, cortical mapping using ESM in a high-density grid is time consuming and becomes impractical due to the sequential pairwise testing of electrodes. In contrast, the mapping of eloquent cortex using the frequency modulation responses from an appropriate cognitive task analyses the signal from all channels simultaneously. Thus, this approach has the potential to significantly improve the general time efficiency of the mapping procedure. Because ESM can cause seizures, after-discharges (Aungaroon et al., 2017; Blume et al., 2004) and, in some cases, pain (e.g., when performed in the proximity parieto-opercular cortex –Mazzola et al., 2012), frequency modulation mapping might also reduce the chance of side effects, potentially allowing the implementation of cortical mapping in high-density ECoG grids. Our procedure to create a probabilistic map could be used to guide ESM mapping in a high-density grid such that boundaries between eloquent and non-eloquent cortex could be identified with ECoG and confirmed with ESM.

Although the DMTS task was not explicitly designed to test whether a specific electrode was eloquent or not, it is fascinating that these data were able to be used to identify the eloquent electrodes with performance similar to that reported in previous studies using tasks specifically designed for that purpose. This aspect points to the wide cortical network that is recruited even when performing a relatively simple task. The used DMTS task involves auditory processing of the syllable, maintenance of the auditory stimulus in short term memory, visual perception of the written cue, comparison of the auditory and visual stimuli for a match to sample decision, a motor response, and error monitoring post-response. It hence engages different relevant stages in human cognition. Nevertheless, we do not argue that the DMTS task used here is the optimal task. An optimal task would presumably include a wider range of motor actions (our task only included moving the index and middle finger of the right hand), a wider range of visual stimuli (to activate e.g. face-place- and motion-sensitive areas), and a language production section – and has to be feasible in the limited amount of time the patient is available for functional cortical mapping. Our main finding is a first proof of concept that the inclusion of low frequency bands, especially the beta band, improves the identification of eloquent electrodes.

We would expect that a similar analysis in a dataset acquired during the performance of a structured cognitive task explicitly designed to activate eloquent cortex would result in better performance of the ECoG mapping. Such tasks should contain language comprehension and production components to activate cortical areas belonging to the language network, different memory components to activate cortical areas of the memory network, and specific motor components, among others. These tasks could be selected according to the probable location of the seizure onset zone and the ECoG implantation scheme. Moreover, similar to the DMTS task used in our study, one task can encompass many cognitive and motor components, allowing the test of multiple functions per task. The use of those different specific tasks could increase the sensitivity and specificity by including the power modulation in the beta band. However, this also represents an increase in the complexity and time require for cognitive testing, thus the task used in the present study seems to be a very good compromise as it tests different cognitive aspects in one relatively simple, easy and short task.

The possibility also exists that including low frequency power was only useful in the context of a non-optimal task. It may be that, in an optimal task all eloquent cortices may be sufficiently activated, such that they can be readily identified using power modulation responses in the broadband gamma alone. Our finding that the beta band was more useful in identifying receptive areas than expressive areas argues against this possibility, since the task was arguably better tuned for identifying receptive areas than expressive areas (e.g. auditory cortex was adequately activated, while IFG (Broca’s area) was only weakly involved during covert rehearsal and reading). Ultimately, additional data or re-analysis of data collected by other groups will be required to clarify this aspect.

### 4.1. Limitations

#### 4.1.1. True gold standard

In order to better understand the validity of cortical mapping with ECoG from cognitive tasks it is relevant to evaluate whether false positives identified by ECoG mapping are in fact false positives or perhaps may be false negatives in the ESM mapping. The true test of whether an area is eloquent or not is the effect of surgical resection of that area on the neuropsychological function after resection. Currently, ESM is the best predictor of the effect of resection on cognitive performance, however to control for the possibility of false negatives in ESM, it will be important to evaluate the cognitive performance of patients after surgical resection. However, these data is not available in our study.

#### 4.1.2. Extension of GLM method to include more indicators

In our analysis we used the power of broadband gamma, alpha and beta bands as indicators for the eloquence of electrodes. However, with the cross-validation procedure implemented in our analysis it is possible in principle to include a multitude of indicators. Other factors that might improve mapping could include power modulations in other frequency bands, power modulations in additional, more specific tasks, and responses measured in other modalities (e.g., functional mapping using fMRI based Blood Oxygenation Level Dependant–BOLD–responses of Transcranial Magnetic Stimulation). Prior predictions about the likelihood of eloquence at a particular cortical area and seizure onset zone (e.g., Positron emission tomography– PET– or EEG) may also be of interest and help to increase mapping accuracy.

#### 4.1.3. From diagnostic ability to diagnostic test

Given the relatively small number of patients included in the present study and the task used, which was developed to study language processing rather than cortical mapping, the method described is not ready to be used as a diagnostic test. In order to further develop our findings into a reliable diagnostic test it will be important to replicate our analysis in a larger population and to compare the results with the presence of neurological or cognitive impairments after neurosurgical resection. With additional data, optimal settings for the test data (frequency bands, time window of interest, task, etc.) and an optimal diagnostic criterion might be identifiable. However, inspection of the probabilistic maps of individual patients in this sample seem to argue against this possibility. They suggest that between-patient variability in the overall level of ECoG responsiveness is too huge to instigate a universal criterion. The optimized ECoG test should be tailored to the individual frequency profile. This then should clearly outperform ESM in predicting presence of neurological or cognitive impairments after neurosurgical resection - before replacing ESM as diagnostic procedure. More likely, the optimized ECoG test will be of use in addition to or as a screening before ESM.

### 4.2. Conclusion

Using advanced signal analysis (time frequency analysis, GLM, AUROC and bootstrapping) in combination to functional and attentional specific analysis we have shown that including alpha and especially beta power modulations from a DMTS task can improve the diagnostic ability in the identification of eloquent cortical areas over the use of broadband gamma alone. We conclude that ECoG mapping may be a useful additional tool to identify eloquent cortical areas but does not replace the need for electrical stimulation mapping. Further studies using tasks specifically designed for eloquent cortical area identification, the use of tailored individual frequency bands, and comparison of different models with post-resection outcomes will elucidate whether this approach could replace or enhance ESM in clinical settings

## Acknowledgments

This work was supported by grants from the Colombian Administrative Department of Science, Technology and Innovation, COLCIENCIAS Colombia (scholarship call 568 to MEAM). The authors would like to thank Dr. S. ten Oever, Dr. J. Peters and Dr. J. Reithler for assistance with data collection during cognitive testing in 2 patients.

## Abbreviations

DMTS: Delayed match-to-sample
ESM: Electrical Stimulation Mapping
ROC: Receiver Operator Characteristic curve
AUROC: Area Under the Receiving Operating Characteristic curve
GLM: Generalized Linear Model

## References

Aoki, F., Fetz, E.E., Shupe, L., Lettich, E., Ojemann, G.A., 1999. Increased gamma-range activity in human sensorimotor cortex during performance of visuomotor tasks. Clin Neurophysiol 110, 524–537.

Arya, R., Horn, P.S., Crone, N.E., 2018. ECoG high-gamma modulation versus electrical stimulation for presurgical language mapping. Epilepsy Behav 79, 26–33. https://doi.org/10.1016/j.yebeh.2017.10.044

Arya, R., Wilson, J.A., Fujiwara, H., Rozhkov, L., Leach, J.L., Byars, A.W., Greiner, H.M., Vannest, J., Buroker, J., Milsap, G., Ervin, B., Minai, A., Horn, P.S., Holland, K.D., Mangano, F.T., Crone, N.E., Rose, D.F., 2017. Presurgical language localization with visual naming associated ECoG high-gamma modulation in pediatric drug-resistant epilepsy. Epilepsia 58, 663–673. https://doi.org/10.1111/epi.13708

Arya, R., Wilson, J.A., Vannest, J., Byars, A.W., Greiner, H.M., Buroker, J., Fujiwara, H., Mangano, F.T., Holland, K.D., Horn, P.S., Crone, N.E., Rose, D.F., 2015. Electrocorticographic language mapping in children by high-gamma synchronization during spontaneous conversation: comparison with conventional electrical cortical stimulation. Epilepsy Res. 110, 78–87. https://doi.org/10.1016/j.eplepsyres.2014.11.013

Aungaroon, G., Zea Vera, A., Horn, P.S., Byars, A.W., Greiner, H.M., Tenney, J.R., Arthur, T.M., Crone, N.E., Holland, K.D., Mangano, F.T., Arya, R., 2017. After-discharges and seizures during pediatric extra-operative electrical cortical stimulation functional brain mapping: Incidence, thresholds, and determinants. Clin Neurophysiol 128, 2078–2086. https://doi.org/10.1016/j.clinph.2017.06.259

Bauer, P.R., Vansteensel, M.J., Bleichner, M.G., Hermes, D., Ferrier, C.H., Aarnoutse, E.J., Ramsey, N.F., 2013. Mismatch between electrocortical stimulation and electrocorticography frequency mapping of language. Brain Stimul 6, 524–531. https://doi.org/10.1016/j.brs.2013.01.001

Blume, W.T., Jones, D.C., Pathak, P., 2004. Properties of after-discharges from cortical electrical stimulation in focal epilepsies. Clin Neurophysiol 115, 982–989. https://doi.org/10.1016/j.clinph.2003.11.023

Bonaiuto, J.J., Meyer, S.S., Little, S., Rossiter, H., Callaghan, M.F., Dick, F., Barnes, G.R., Bestmann, S., 2018. Lamina-specific cortical dynamics in human visual and sensorimotor cortices. Elife 7. https://doi.org/10.7554/eLife.33977

Bouchard, K.E., Mesgarani, N., Johnson, K., Chang, E.F., 2013. Functional organization of human sensorimotor cortex for speech articulation. Nature 495, 327–332. https://doi.org/10.1038/nature11911

Brosch, M., Budinger, E., Scheich, H., 2002. Stimulus-related gamma oscillations in primate auditory cortex. J. Neurophysiol. 87, 2715–2725. https://doi.org/10.1152/jn.2002.87.6.2715

Brunner, P., Ritaccio, A.L., Lynch, T.M., Emrich, J.F., Wilson, J.A., Williams, J.C., Aarnoutse, E.J., Ramsey, N.F., Leuthardt, E.C., Bischof, H., Schalk, G., 2009. A practical procedure for real-time functional mapping of eloquent cortex using electrocorticographic signals in humans. Epilepsy & Behavior 15, 278–286. https://doi.org/10.1016/j.yebeh.2009.04.001

Buzsaki, G., 2004. Neuronal Oscillations in Cortical Networks. Science 304, 1926–1929. https://doi.org/10.1126/science.1099745

Buzsáki, G., Anastassiou, C. a., Koch, C., 2012. The origin of extracellular fields and currents — EEG, ECoG, LFP and spikes. Nat. Rev. Neurosci. 13, 407–420. https://doi.org/10.1038/nrn3241

Crone, N.E., Miglioretti, D.L., Gordon, B., Lesser, R.P., 1998a. Functional mapping of human sensorimotor cortex with electrocorticographic spectral analysis. II. Event-related synchronization in the gamma band. Brain 121 (Pt 12), 2301–2315.

Crone, N.E., Miglioretti, D.L., Gordon, B., Sieracki, J.M., Wilson, M.T., Uematsu, S., Lesser, R.P., 1998b. Functional mapping of human sensorimotor cortex with electrocorticographic spectral analysis. I. Alpha and beta event-related desynchronization. Brain 121 (Pt 12), 2271–2299.

Crone, N.E., Sinai, A., Korzeniewska, A., 2006. High-frequency gamma oscillations and human brain mapping with electrocorticography. Prog. Brain Res. 159, 275–295. https://doi.org/10.1016/S0079-6123(06)59019-3

de Pesters, A., Coon, W.G., Brunner, P., Gunduz, A., Ritaccio, A.L., Brunet, N.M., de Weerd, P., Roberts, M.J., Oostenveld, R., Fries, P., Schalk, G., 2016. Alpha power indexes task-related networks on large and small scales: A multimodal ECoG study in humans and a non-human primate. NeuroImage 134, 122–131. https://doi.org/10.1016/j.neuroimage.2016.03.074

Escabí, M.A., Read, H.L., Viventi, J., Kim, D.-H., Higgins, N.C., Storace, D.A., Liu, A.S.K., Gifford, A.M., Burke, J.F., Campisi, M., Kim, Y.-S., Avrin, A.E., Spiegel Jan, V. der, Huang, Y., Li, M., Wu, J., Rogers, J.A., Litt, B., Cohen, Y.E., 2014. A high-density, high-channel count, multiplexed μECoG array for auditory-cortex recordings. J. Neurophysiol. 112, 1566–1583. https://doi.org/10.1152/jn.00179.2013

Felleman, D.J., Van Essen, D.C., 1991. Distributed hierarchical processing in the primate cerebral cortex. Cerebral cortex (New York, N.Y. : 1991) 1, 1–47. https://doi.org/10.1093/cercor/1.1.1

Fries, P., Nikolić, D., Singer, W., 2007. The gamma cycle. Trends in Neurosciences 30, 309–316. https://doi.org/10.1016/j.tins.2007.05.005

Gray, C.M., König, P., Engel, A.K., Singer, W., 1989. Oscillatory responses in cat visual cortex exhibit inter-columnar synchronization which reflects global stimulus properties. Nature 338, 334–337. https://doi.org/10.1038/338334a0

Green, D.M., Swets, J.A., 2000. Signal detection theory and psychophysics, Repr. ed. ed. Peninsula Publ, Los Altos Hills, Calif.

Hamberger, M.J., 2007. Cortical language mapping in epilepsy: A critical review. Neuropsychology Review 17, 477–489. https://doi.org/10.1007/s11065-007-9046-6

Hamilton, L.S., Edwards, E., Chang, E.F., 2018. A spatial map of onset and sustained responses to speech in human superior temporal gyrus. Current Biology (in press), 1–12. https://doi.org/10.1016/j.cub.2018.04.033

Hermes, D., Miller, K.J., Vansteensel, M.J., Aarnoutse, E.J., Leijten, F.S.S., Ramsey, N.F., 2012. Neurophysiologic correlates of fMRI in human motor cortex. Hum Brain Mapp 33, 1689–1699. https://doi.org/10.1002/hbm.21314

Hermiz, J., Rogers, N., Kaestner, E., Ganji, M., Cleary, D.R., Carter, B.S., Barba, D., Dayeh, S.A., Halgren, E., Gilja, V., 2018. Sub-millimeter ECoG pitch in human enables higher fidelity cognitive neural state estimation. Neuroimage 176, 454–464. https://doi.org/10.1016/j.neuroimage.2018.04.027

Jensen, O., Mazaheri, A., 2010. Shaping Functional Architecture by Oscillatory Alpha Activity: Gating by Inhibition. Frontiers in Human Neuroscience 4, 1–8. https://doi.org/10.3389/fnhum.2010.00186

Kundu, S., Kers, J.G., Janssens, A.C.J.W., 2016. Constructing Hypothetical Risk Data from the Area under the ROC Curve: Modelling Distributions of Polygenic Risk. PLoS ONE 11, e0152359. https://doi.org/10.1371/journal.pone.0152359

Lachaux, J.-P., Jerbi, K., Bertrand, O., Minotti, L., Hoffmann, D., Schoendorff, B., Kahane, P., 2007. A blueprint for real-time functional mapping via human intracranial recordings. PLoS ONE 2, e1094. https://doi.org/10.1371/journal.pone.0001094

Lachaux, J.P., Rudrauf, D., Kahane, P., 2003. Intracranial EEG and human brain mapping. J. Physiol. Paris 97, 613–628. https://doi.org/10.1016/j.jphysparis.2004.01.018

Lee, H.W., Webber, W.R.S., Crone, N., Miglioretti, D.L., Lesser, R.P., 2010. When Is Electrical Cortical Stimulation More Likely To Produce Afterdischarges? Clin Neurophysiol 121, 14–20. https://doi.org/10.1016/j.clinph.2009.10.001

Lesser, R.P., Lüders, H., Klem, G., Dinner, D.S., Morris, H.H., Hahn, J., 1985. Ipsilateral trigeminal sensory responses to cortical stimulation by subdural electrodes. Neurology 35, 1760–1763.

Lesser, R.P., Lüders, H., Klem, G., Dinner, D.S., Morris, H.H., Hahn, J., 1984. Cortical afterdischarge and functional response thresholds: results of extraoperative testing. Epilepsia 25, 615–621.

Leszczynski, M., Barczak, A., Kajikawa, Y., Ulbert, I., Falchier, A., Tal, I., Haegens, S., Melloni, L., Knight, R., Schroeder, C., 2019. Dissociation of broadband high-frequency activity and neuronal firing in the neocortex: Supplementary Materials (preprint). Neuroscience. https://doi.org/10.1101/531368

Leuthardt, E.C., Miller, K., Anderson, N.R., Schalk, G., Dowling, J., Miller, J., Moran, D.W., Ojemann, J.G., 2007. Electrocorticographic frequency alteration mapping: a clinical technique for mapping the motor cortex. Neurosurgery 60, 260–270; discussion 270-271. https://doi.org/10.1227/01.NEU.0000255413.70807.6E

Mazzola, L., Isnard, J., Peyron, R., Mauguière, F., 2012. Stimulation of the human cortex and the experience of pain: Wilder Penfield’s observations revisited. Brain 135, 631–640. https://doi.org/10.1093/brain/awr265

McCullagh, P., Nelder, J.A., 1998. Generalized linear models, 2nd ed. ed, Monographs on statistics and applied probability. Chapman & Hall/CRC, Boca Raton.

Mesgarani, N., Cheung, C., Johnson, K., Chang, E.F., 2014. Phonetic feature encoding in human superior temporal gyrus. Science 343, 1006–1010. https://doi.org/10.1126/science.1245994

Miller, K.J., Leuthardt, E.C., Schalk, G., Rao, R.P.N., Anderson, N.R., Moran, D.W., Miller, J.W., Ojemann, J.G., 2007. Spectral changes in cortical surface potentials during motor movement. J. Neurosci. 27, 2424–2432. https://doi.org/10.1523/JNEUROSCI.3886-06.2007

Mooij, A.H., Sterkman, L.C.M., Zijlmans, M., Huiskamp, G.J.M., 2018. Electrocorticographic high gamma language mapping: Mind the pitfalls of comparison with electrocortical stimulation. Epilepsy Behav 82, 196–199. https://doi.org/10.1016/j.yebeh.2018.02.001

Muller, L., Hamilton, L.S., Edwards, E., Bouchard, K.E., Chang, E.F., 2016. Spatial resolution dependence on spectral frequency in human speech cortex electrocorticography. J Neural Eng 13, 056013. https://doi.org/10.1088/1741-2560/13/5/056013

Murthy, V.N., Fetz, E.E., 1992. Coherent 25- to 35-Hz oscillations in the sensorimotor cortex of awake behaving monkeys. Proceedings of the National Academy of Sciences 89, 5670–5674. https://doi.org/10.1073/pnas.89.12.5670

Nagasawa, T., Rothermel, R., Juhász, C., Fukuda, M., Nishida, M., Akiyama, T., Sood, S., Asano, E., 2010. Cortical gamma-oscillations modulated by auditory-motor tasks-intracranial recording in patients with epilepsy. Hum Brain Mapp 31, 1627–1642. https://doi.org/10.1002/hbm.20963

Nelder, J.A., Wedderburn, R.W.M., 1972. Generalized Linear Models. Journal of the Royal Statistical Society. Series A (General) 135, 370. https://doi.org/10.2307/2344614

Ojemann, G., Ojemann, J., Lettich, E., Berger, M., 1989. Cortical language localization in left, dominant hemisphere. Journal of Neurosurgery 71, 316–326. https://doi.org/10.3171/jns.1989.71.3.0316

Oostenveld, R., Fries, P., Maris, E., Schoffelen, J.M., 2011. FieldTrip: Open source software for advanced analysis of MEG, EEG, and invasive electrophysiological data. Computational Intelligence and Neuroscience 2011. https://doi.org/10.1155/2011/156869

Penfield, W., Boldrey, E., 1937. Somatic Motor and Sensory Representation in Man. Brain 389–443. https://doi.org/10.1093/brain/60.4.389

Pfurtscheller, G., Neuper, C., Kalcher, J., 1993. 40-Hz oscillations during motor behavior in man. Neurosci. Lett. 164, 179–182.

Scheeringa, R., Fries, P., 2017. Cortical layers, rhythms and BOLD signals. Neuroimage. https://doi.org/10.1016/j.neuroimage.2017.11.002

Sinai, A., Bowers, C.W., Crainiceanu, C.M., Boatman, D., Gordon, B., Lesser, R.P., Lenz, F.A., Crone, N.E., 2005. Electrocorticographic high gamma activity versus electrical cortical stimulation mapping of naming. Brain 128, 1556–1570. https://doi.org/10.1093/brain/awh491

Vansteensel, M.J., Bleichner, M.G., Dintzner, L.T., Aarnoutse, E.J., Leijten, F.S.S., Hermes, D., Ramsey, N.F., 2013. Task-free electrocorticography frequency mapping of the motor cortex. Clin Neurophysiol 124, 1169–1174. https://doi.org/10.1016/j.clinph.2012.08.048

Vansteensel, M.J., Hermes, D., Aarnoutse, E.J., Bleichner, M.G., Schalk, G., van Rijen, P.C., Leijten, F.S.S., Ramsey, N.F., 2010. Brain-computer interfacing based on cognitive control. Annals of Neurology n/a-n/a. https://doi.org/10.1002/ana.21985

Wada, J., Rasmussen, T., 2007. Intracarotid injection of sodium amytal for the lateralization of cerebral speech dominance. 1960. J. Neurosurg. 106, 1117–1133. https://doi.org/10.3171/jns.2007.106.6.1117

Wang, Y., Fifer, M.S., Flinker, A., Korzeniewska, A., Cervenka, M.C., Anderson, W.S., Boatman-Reich, D.F., Crone, N.E., 2016. Spatial-temporal functional mapping of language at the bedside with electrocorticography. Neurology 86, 1181–1189. https://doi.org/10.1212/WNL.0000000000002525

Wu, M., Wisneski, K., Schalk, G., Sharma, M., Roland, J., Breshears, J., Gaona, C., Leuthardt, E.C., 2010. Electrocorticographic Frequency Alteration Mapping for Extraoperative Localization of Speech Cortex. Neurosurgery 66, E407–E409. https://doi.org/10.1227/01.NEU.0000345352.13696.6F

